# Extracellular matrix dependent regulation of Septin 7 in focal adhesions promotes mechanosensing and response in fibroblasts

**DOI:** 10.1101/2024.02.27.582035

**Authors:** Wesley Sturgess, Swathi Packirisamy, Rodina Geneidy, Vinay Swaminathan

## Abstract

Fibroblasts are contractile adherent cells that maintain tissue homeostasis by sensing a wide array of changes in the extracellular matrix (ECM) and in response, regulate the physical and compositional properties of the ECM. These diverse cues are sensed by focal adhesions (FAs) that differentially couple changes in the ECM to the actomyosin machinery via modulation of integrin activation and the resultant recruitment of several proteins. One such protein is Septin-7 (Sept-7) that belongs to the septin family and has been found in FA proteomics and interactome studies. Sept-7 however, is not considered an FA protein and is thought to regulate and be regulated by actin outside of FAs. To reconcile these differences, here we used total internal reflection microscopy to image Sept-7 localization and dynamics at the cell-ECM interface and found that that ECM-mediated integrin activation in fibroblasts regulates the formation of spatially distinct higher order Sept-7 structures at FA subpopulations. In and around FAs located in the perinuclear regions of the cell, ECM binding resulted in the formation and stabilization of Sept-7 bundles while ECM binding and complete integrin activation promoted the growth of FA-like elongated Sept-7 structures that dynamically associated with the core of peripheral FAs. Functionally, peripheral Sept-7 structures promoted the elongation of peripheral FAs while perinuclear Sept-7 bundles were critical in regulating the maturation and stabilization of perinuclear FAs. Due to this coupling between the ECM, integrin activation and regulation of Sept-7 structures, we found that Sept-7 is required for a wide range of ECM sensing functions in fibroblasts including modulating sensitivity to changes in ECM stiffness and density and in contributing to the cells ability to remodel the ECM. Collectively, our results show that Sept-7 is an FA protein that gets recruited and assembled in diverse higher order structures in an ECM dependent manner to differentially regulate FA subpopulations and promote mechanosensing and ECM remodelling functions in fibroblasts.

## Introduction

The extracellular matrix (ECM) is a complex, multicomponent and dynamically changing non-cellular structure surrounding most cell types and tissues in our body. The proper regulation of the ECM’s physical and biochemical properties is critical for several cellular processes, from cell specification and organogenesis to wound healing and immune responses and is altered in diseases such as cancer and atherosclerosis^1–6^. Fibroblasts are one of the primary cell types responsible for this regulatory role and do so by sensing changes in ECM properties and responding to these changes by producing, modifying, and remodelling the ECM^7^. Fibroblasts thus need to have highly sensitive ECM-sensing mechanisms that allow it to sense small and distinct changes in the ECM environment that downstream trigger highly specific cellular responses. This important function is primarily mediated by integrin-based focal adhesions (FAs)^8–11^.

FAs comprise of the integrin family of receptors that indirectly couples the ECM to the cytoskeleton via a network of proteins called the adhesome. The complex regulation of recruitment and activity of adhesome proteins coupled to ECM-dependent integrin activation modulates the biophysical coupling between the ECM and the cytoskeleton and drives the sensing of different ECM cues as well as regulating downstream cellular responses^12^. While we now know of several mechanisms by which FAs can sense changes in ECM stiffness and architecture, mechanisms that fine tune the sensitivity and specificity of ECM sensing and cellular response is still not completely known. Proteomics using different FA isolation techniques and across different cell types have identified more than 2000 proteins in the adhesome^13–15^, and this has led to the hypothesis that many of these proteins could be responsible for this sensitivity and specificity. However, currently several of these proteins have no identified function in regulating FAs or in ECM sensing and cellular response. Additionally, since most of the proteomics studies rely on bulk isolation of FAs in the cell, information about composition of FA subpopulations within a cell based on location, maturation level or other subcellular states and whether these subpopulations play specific roles is not known.

To address these specific questions, we focussed on investigating the role of Sept-7 which is one of the most enriched septins in the adhesome^15^. Septins are GTP-binding proteins that self-assemble into oligomers and polymers and form higher ordered structures either with linear or with curved filaments and rings^16^. Several studies have identified important roles for septins in regulating FA formation, maturation and disassembly^17,18^. In addition, Sept-7 containing bundles are found in proximity to FAs at the cell periphery^17^ as well as in the perinuclear area along actin fibers where through interactions with F-actin, Sept-7 plays an important role in sensing of ECM stiffness^19^. However, in spite of Sept-7 being found in FA proteomic studies and evidence suggesting interactions between Sept-7 and other FA proteins, septins and specifically Sept-7 is considered to be excluded from FAs and its direct relationship with changing ECM cues and integrin activation is not known^20–22^.

We aimed to resolve these differences and understand the relationship between ECM sensing, integrin activation and of Sept-7 in FAs in this study. By using total internal reflection microscopy (TIRFM) to image the cell-ECM interface with high resolution as well as biochemical purification of adhesion complexes using ECM-coated beads, we found that Sept-7 forms distinct higher order structures that localize to FAs of mouse embryonic fibroblasts (MEFs). In addition to location specific distinct high order architecture, Sept-7 localization was also spatially and temporally distinctly localized within FA subpopulations. Sept-7 recruitment to the back of perinuclear FAs (in close proximity to the nucleus) and formation of higher order bundles was dependent on binding to the ECM protein fibronectin(FN) while FN binding and complete integrin activation was required to form elongated FA-like Sept-7 structures within the core of peripheral FAs (closer to the leading edge). To test the function of Sept-7 in FA regulation, we downregulated Sept-7 expression in MEFs and found that this led to a dramatic loss in the perinuclear FA population by affecting perinuclear FA maturation rate and lifetime, but Sept-7 loss had only a minor effect on peripheral FA elongation. The loss of perinuclear FAs due to downregulation of Sept-7 correlated with significant loss of sensitivity to changes in FN density in MEFs as well as their ability to remodel and clear the FN. Taken together, our work identifies Sept-7 as an adhesome protein that via ECM dependent changes in its higher order architecture and localization regulates FA populations. Through these mechanisms, Sept-7 promotes the sensitivity of fibroblasts to regulate the physical properties of the ECM.

## Results

### SEPT-7 localizes in spatially and temporally distinct patterns in FA sub-populations

Based on microscopy-based localization data, Sept-7 is thought to be excluded from FAs which contradicts proteomics studies where Sept-7 is found to be enriched in FAs^15,23,24^. Since FA composition and function is highly dependent on the cell type, we sought to clarify these differences using fibroblasts which are highly contractile adherent cells that form large, dynamic FAs for motility and ECM sensing. To test if Sept-7 is associated with FAs, we first utilized an ECM coated magnetic bead-based assay that allows for isolation of the adhesion fraction and probed for proteins of interests using western blotting^25–27^. Briefly, fibronectin-coated beads were added to mouse embryonic fibroblasts (MEFs), incubated for 30 minutes, and then lysed (Figure S1A). The bead-bound fraction was then isolated, probed and compared to the total lysate. Blotting for the FA proteins vinculin, talin, paxillin and FAK as well as for tyrosine phosphorylated FAK and paxillin showed enrichment of these proteins in the bead fraction, while GAPDH was only present in the total lysate and absent in the bead fraction (Figure S1B). This verified our methodology and confirmed that the bead fraction was indeed the adhesion fraction. We then probed this adhesion fraction for Sept-7 along with the other FA proteins and found that Sept-7 is also significantly enriched in the adhesion fraction (Figure S1B). This suggests that Sept-7 is indeed associated with canonical FA proteins and is present in adhesions formed on ECM-coated beads.

We then sought to determine if Sept-7 also localizes to FAs formed at the cell-ECM substrate interface. We plated MEFs on glass-bottom dishes coated with 10 µg/ml fibronectin (FN) which allows for formation of robust integrin-dependent FAs that promote optimal ECM sensing for cell migration and mechanotransduction^28^. Cells were then fixed and co-immunostained for the FA protein paxillin, F-actin, and Sept-7 and imaged using TIRFM (Figure 1A). Consistent with previous results, we found filamentous Sept-7 decorating ventral actin stress fibers near paxillin enriched FAs beneath and around the nucleus (Figure 1B, top panel)^16^. In addition, we also observed Sept-7 near the front of the cell in the lamellipodia as well as in FA-shaped structures in the lamella of the cell where it seemed to co-localize with the paxillin signal (Figure 1B, lower panel). The lamellipodia and lamella-localized Sept-7 signal while robust however was relatively weak compared to the signal from the filamentous structures found in proximity of the nucleus. Due to the 2 distinct structures of Sept-7 observed in the proximity of FAs, and to determine the exact location of Sept-7 relative to F-actin and paxillin, we first classified the paxillin stained FAs as either perinuclear (connected to ventral actin stress fibers beneath the nucleus) or peripheral (located closer to the edge of the cell). We then generated a series of line scans across the FAs and plotted the average location of Sept-7 and F-actin relative to the location of paxillin within the FA (Figure 1C). This analysis showed that in perinuclear FAs, filamentous Sept-7 localized towards the rear of the FA with its average intensity peak outside of the FA and a partial overlap with paxillin. In contrast, at the peripheral FAs, puncta or elongated FA-like structures of Sept-7 peaked more centrally to both the paxillin and the F-actin peak and terminated at the end of the FA (Figure 1C).

**Figure 1.**
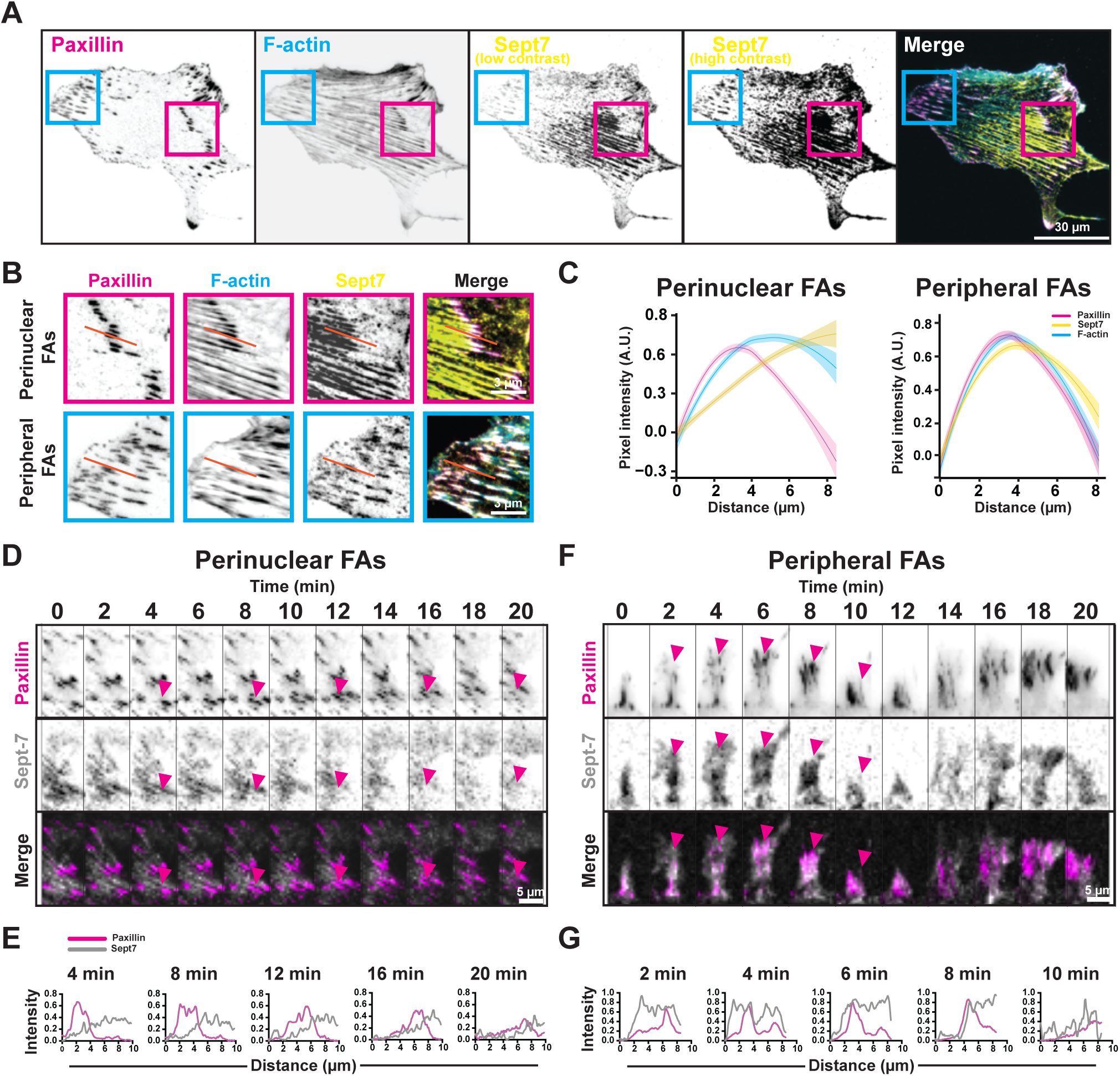
Sept-7 localizes in spatially and temporally distinct patterns in FA sub-populations. (A) Representative TIRFM images of MEFs stained with paxillin (magenta), F-actin (cyan), and Sept-7, (yellow, low, and high contrast). Magenta and cyan boxes highlight perinuclear and peripheral FAs respectively. (B) Insets of perinuclear (magenta), and peripheral (cyan) FAs from (A), red lines represent example FA line scans. (C) Curve plots showing normalized line scan intensities of paxillin, Sept-7, and F-actin, for perinuclear(left) and peripheral(right) FAs, shaded curves represent SD, (n = 5 cells, 10–12 line scans per adhesion type). (D and F) Montages taken from live TIRFM movie, of perinuclear and peripheral FAs from MEFs co-transfected with mCherry-paxillin (magenta, top) and YFP-Sept-7 (grey, middle), and merged channels (bottom), magenta arrows highlight FAs over time. (E and G) line plots showing colocalization of paxillin (magenta) and Sept-7 (grey) at FAs overtime indicated by arrows in (D and F).

This FA subpopulation-dependent localization and distinct higher order structures of Sept-7 suggested to us that the dynamics of Sept-7 in these populations could also be differentially coupled to these FA subpopulations. To investigate this, we co-expressed Sept-7-YFP with paxillin-mCherry in MEFs and imaged cells 4 hours after plating them on 10 µg/ml FN coated glass-bottom dishes using TIRFM (Figure 1D and F, supplementary movies (SM1)). Time-lapse imaging confirmed distinct dynamics of localization at the peripheral FAs compared to perinuclear FAs. In perinuclear FAs, Sept-7 bundles localized to the back of the FA and remained stably associated during the entire lifetime of the perinuclear FA (Figure 1E). The time-period of localization of Sept-7 was coupled to the lifetime of the perinuclear FAs which were relatively long (at least > 10 minutes). In contrast, Sept-7 puncta appeared along with newly formed peripheral FAs and followed the fate of the FAs, with puncta disappearing with FAs turning over or remaining stably associated with the FAs and maturing into larger structures in the lamellar region of the cell (Figure 1G). Taken together, our data here shows that Sept-7 robustly localizes to FAs of adherent cells in distinct higher order structures with dynamics and FA localization dependent on the FA subpopulation within the cell.

### ECM-mediated integrin activation promotes formation of higher order Sept-7 structures and its association with FAs

Our data on Sept-7 localization and the coupling of its dynamics to FA dynamics suggests a mechanistic link between Sept-7 recruitment to FAs, formation of higher order structures and integrin activation which regulates the formation and fate of FAs. To investigate this, we plated MEFs on glass-bottom dishes coated with poly-l-lysine (PLL-to prevent ECM mediated integrin activation), or 0.1 µg/ml FN (to achieve low levels of integrin activation) and fixed and stained the cells for Sept-7 and paxillin to compare with cells on 10 µg/ml FN (Figure 2A). As expected, cells coated on PLL had no large perinuclear or peripheral FAs as stained by paxillin compared to cells on 10 µg/ml FN, which was further confirmed by quantification of FA size which showed an expected ECM density dependent increase (Figure S2A, S2B). The loss of FAs on cells plated on PLL also coincided with loss of filamentous Sept-7 structures in the perinuclear region with Sept-7 instead forming puncta or rings throughout the cell (Figure 2A). Increasing the FN concentration to 0.1µg/ml FN resulted in formation of perinuclear Sept-7 bundles though peripheral Sept-7 structures were still punctate-like unlike cells on 10 µg/ml FN with more elongated structures (Figure 2A). To quantify these morphologies of Sept-7 structures, we measured Sept-7 co-alignment or anisotropy for the perinuclear bundles and the length and shape of peripheral Sept-7 structures across these conditions (Figure 2B, S2C). This quantification showed a robust increase in perinuclear Sept-7 anisotropy which coincided with increase in perinuclear FA size upon ECM binding (Figure 2B, S2B). Peripheral Sept-7 structures on the other hand, only started getting elongated at the highest ECM density of 10µg/ml FN with no statistical difference between PLL and 0.1µg/ml FN (Figure 2B, S2C) which correlated with smaller peripheral FA size in 0.1µg/ml FN compared to 10µg/ml FN (Figure S2B). This suggested to us that the elongation of peripheral Sept-7 structures could be actomyosin driven since myosin II contractility in MEFs drives FA growth^29^. Unsurprisingly, inhibiting contractility using blebbistatin on MEFs plated on 10µg/ml FN or plating cells on polyacrylamide (PAA) gels of low rigidity (0.4KPa) resulted in loss of elongated peripheral Sept-7 structures as well as perinuclear Sept-7 bundles (Figure S2D). Thus, formation of higher order Sept-7 structures correlates with FA morphology and growth which is sensitive to changes in ECM binding, stiffness and actomyosin contractility.

**Figure 2.**
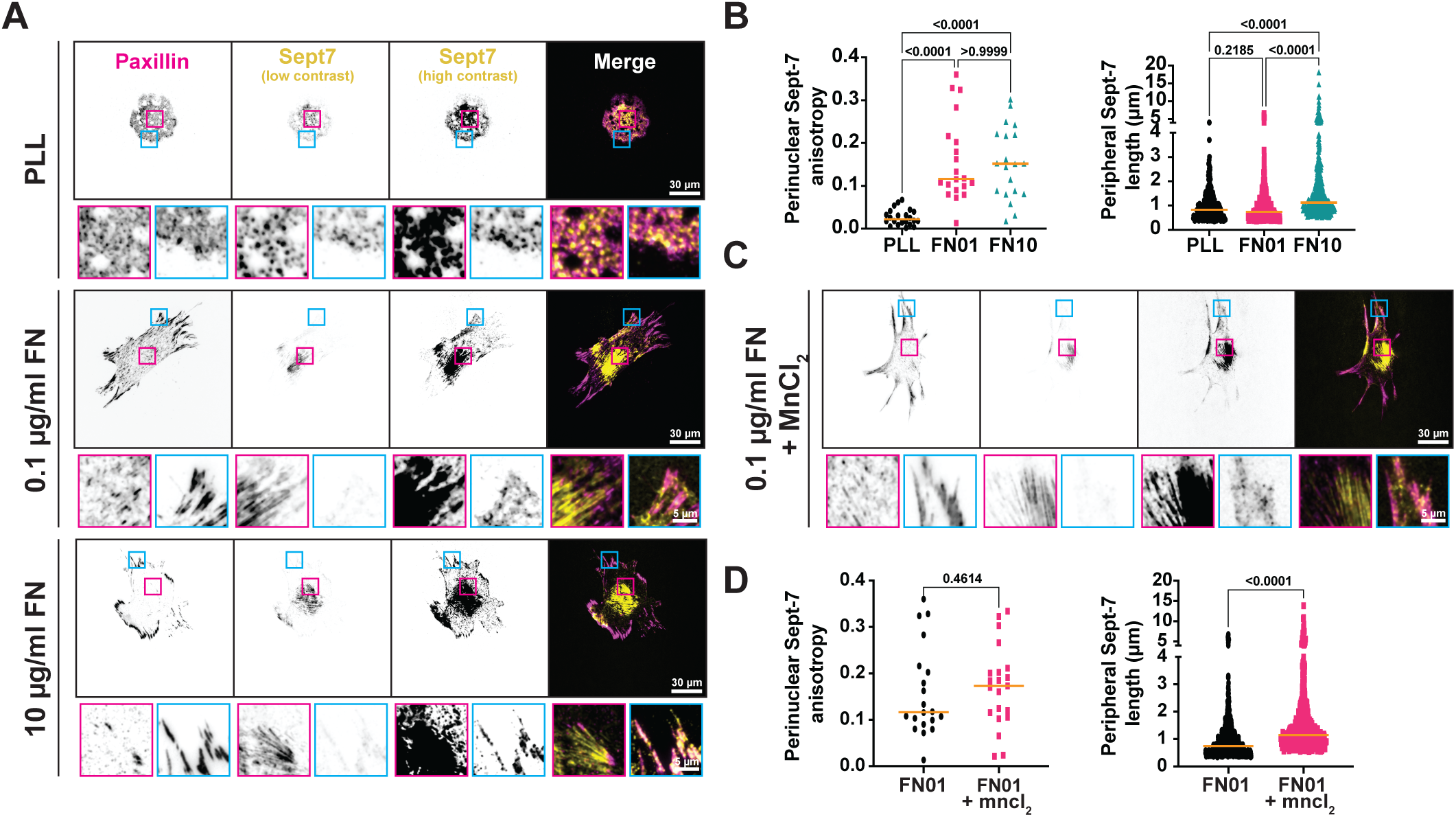
ECM-mediated Integrin activation promotes formation of filamentous perinuclear Sept-7 and its association with FAs. (A) TIRFM images showing paxillin (magenta), Sept-7 (yellow, low, and high contrasts), and merge in MEFs plated on PLL, 0.1 µg/ml FN, and 10 µg/ml FN, magenta and cyan insets highlight perinuclear and peripheral FAs respectively. (B) Quantification of Sept-7 anisotropy, n = 20 – 22 cells, and Sept-7 peripheral structure major axis length, n = 327 – 683 puncta of MEFs plated on PLL, 0.1 µg/ml FN, and 10 µg/ml FN, with Kruskal-Wallis and Dunns multiple comparisons test. (C) TIRFM images show paxillin (magenta), Sept-7 (yellow, low, and high contrasts), and merge of MEFs plated on 0.1 µg/ml FN + 1mM MnCl2, magenta and cyan insets highlight perinuclear and peripheral FAs respectively. (D) Quantification of Sept-7 anisotropy, n = 20 – 21 cells, and Sept-7 peripheral structure major axis length, n = 527 – 683 puncta of MEFs plated on 0.1 µg/ml FN (from 2B), and 0.1 µg/ml FN + 1mM MnCl2, with Mann-Whitney test.

Next, we wanted to test the specific role of integrin activation in formation of Sept-7 structures. To do so, we pre-treated cells with 1 mM MnCl_2_ to shift surface expressed integrins to an extended (primed) conformation, and then plated them on 0.1µg/ml FN prior to fixing and immunostaining for paxillin and Sept-7 (Figure 2C)^30,31^. We found that treating MEFs with MnCl_2_ resulted in a robust increase in FA size compared to untreated cells thus suggesting an increase in integrin activation at 0.1µg/ml FN (Figure S2E). Analysis of Sept-7 structures showed that this increase in integrin activation resulted in an insignificant increase in perinuclear Sept-7 anisotropy compared to cells on 0.1µg/ml FN (Figure 2D). In peripheral regions however, increased integrin activation on 0.1µg/ml led to a complete restoration of elongated Sept-7 structures to levels of 10 µg/ml FN (Figure 2D). Taken together, these results show that ECM-mediated integrin activation differentially regulates the formation and coupling with FAs of distinct higher order Sept-7 structure on the ventral cell surface.

### Sept-7 promotes the maturation and stabilization of perinuclear FAs

Due to the strong coupling between formation of higher order Sept-7 structures and FAs, we next investigated the role of Sept-7 in FA formation and dynamics. To do this, we used siRNAs to knockdown Sept-7 (Sept-7 KD) expression in MEFs (Figure S3A) and quantified FA morphodynamics using live TIRFM. To measure static FA properties, we fixed Sept-7 KD or non-targeting (NT) siRNA control cells plated on 10µg/ml FN and stained for paxillin and F-actin (Figure 3A). Strikingly, we observed a near complete loss of large perinuclear FAs and associated ventral stress fibers in Sept-7 KD cells compared to NT controls, accompanied with a relatively less apparent effect on peripheral FAs (Figure 3A and B). Quantification of FA number and size confirmed our observations, showing a significant reduction in the number of perinuclear FAs in Sept-7 KD cells compared to NT control (Figure 3C). In addition to the reduced number, Sept-7 loss also resulted in the remaining perinuclear FAs to be significantly smaller compared to the controls (Figure 3D). Knocking down Sept-7 had no effect on the average number of peripheral FAs though quantification revealed a slight reduction in peripheral FA size compared to the NT control (Figure 3C and D). To test if these effects of Sept-7 loss was specific to the ventral surface of the cell, we used 3D Structured Illumination Microscopy (SIM) to image the ventral actin stress fibers which are linked to perinuclear FAs, and the apical perinuclear actin cap which traverse the cell and attach to peripheral FAs (Figure S3C). In NT siRNA control cells, we again found robust actin stress fibers above and below the nucleus which were associated with Sept-7 bundles on the ventral side and smaller Sept-7 structures on the apical side (Figure S3B, left panel). Loss of Sept-7 however only affected the ventral actin stress fibers linked to perinuclear FAs resulting in loss of thick bundles with little or no effect on the apical perinuclear actin cap (Figure S3B, right panel).

**Figure 3.**
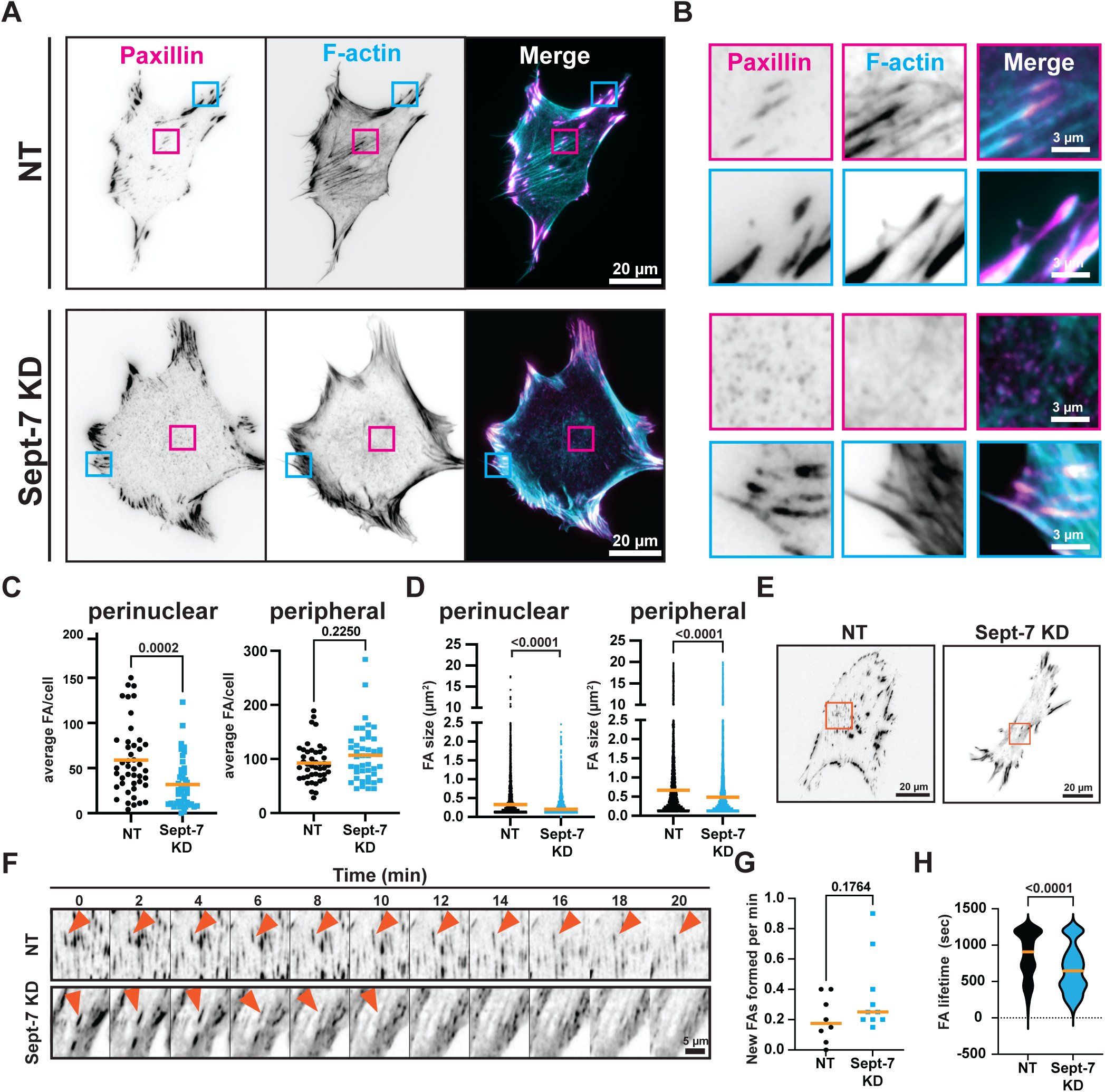
Sept-7 promotes the maturation and stabilization of perinuclear FAs. (A) TIRFM images show MEFs transfected with NT (top panels) or Sept-7 KD (lower panels) siRNA and labelled with paxillin (magenta) and F-actin (cyan), and merge, magenta and cyan boxes highlight perinuclear and peripheral FAs respectively. (B) ROIs of perinuclear (magenta) and peripheral (cyan) FAs from (A). (C) Quantification of average FA count per cell, and (D) FA size for perinuclear and peripheral FA sub-populations n = 44 cells. (E) Live cell TIRFM image showing NT and Sept-7 KD MEFs transfected with mCherry-paxillin, orange boxes highlight perinuclear FA ROIs. (F) Montage of 20 min live cell TIRFM movie of perinuclear FAs of NT control(top) and Sept-7 KD(bottom) cells, orange arrows highlight an example FA over time. (G) Quantification of FA formation rate taken from kymograph quantification of live cell TIRFM videos, n = 4 cells. (H) Quantification of minimum FA lifetime of NT and Sept-7 KD cells, n = 53–75 FAs. All statistic performed using a Mann-Whitney test.

Next, due to the strong effect on perinuclear FAs, we asked if the significant reduction in the number and size of perinuclear FAs accompanying Sept-7 loss was due to reduction in formation of new perinuclear FAs or due to changes in the perinuclear FA lifetime and growth. To answer this, we transfected NT control and Sept-7 KD cells with paxillin-mCherry and imaged cells live using TIRFM (Figure 3E and F, supplementary movie SM2). Examination of timelapse movies revealed that in NT siRNA expressing cells, paxillin-mCherry localized to peripheral and perinuclear FAs and in peripheral FAs showed formation, turnover, and maturation dynamics similar to previous published reports^28^. Unlike peripheral FAs, perinuclear FAs in NT control MEFs were more stable with longer lifetimes with fewer new perinuclear FAs forming during the course of 10-20 minutes of image acquisition. In fact, perinuclear FAs formed prior to starting of acquisition lasted for longer than 10 minutes before disassembling (Figure 3E and F). In contrast, consistent with immunostaining data, in Sept-7 KD MEFs, paxillin-mCherry localized to peripheral FAs but was either completely absent in the perinuclear regions or present only in small perinuclear FA-like structures (Figure 3E, supplementary move SM3). While we couldn’t observe any differences in peripheral FA dynamics within the temporal resolution of our acquisition, the existing perinuclear FAs in Sept-7 KD cells disassembled rapidly compared to NT controls (Figure 3F). In addition, we also observed formation of several perinuclear paxillin puncta during the course of 10-20 minutes of image acquisition that disassembled instead of maturing into larger perinuclear FAs (Figure 3F). To quantify these dynamics, we used kymograph-based analysis and measured perinuclear FA formation rate and its observable average lifetime (Figure 3G and H). This analysis confirmed that while loss of Sept-7 resulted in a slight but not statistically significant increase in the formation rate of small perinuclear paxillin positive puncta, it led to a significant reduction in its minimum lifetime compared to NT controls. Thus, our data shows that ECM-dependent Sept-7 bundles promote the stabilization of perinuclear FAs by increasing the maturation rate and lifetime of perinuclear FAs in adherent cells.

### SEPT-7 enhances sensitivity of cells to changes in ECM cues and contributes to the cell’s ability to remodel the ECM

Our results from above show that physical cues from the ECM can regulate the localization-specific architecture of Sept-7 and that this is differentially mediated through integrin activation. Additionally, we found that this localization-specific architecture of Sept-7 can in turn regulate the stability and dynamics of specific sub-populations of FAs. This led us to hypothesize that Sept-7 is critical for cellular functions that depend on FA-mediated sensing of changes in the ECM which rely on integrin activation. To investigate this, we first tested the role of Sept-7 in sensing changes in ECM rigidity by plating MEFs on PAA gels with stiffness of 0.4KPa and 60KPa and measuring changes in cell area across these conditions (Figure 4A). We first verified that increasing the stiffness from 0.4KPa to 60KPa did indeed result in increase in higher order Sept-7 structures in NT control MEFs which correlated with increase in cell spread area on 60KPa compared to 0.4 KPa PAA gels (Figure 4B, S3D). However, while Sept-7 KD MEFs spread to the same size on 0.4KPa gels compared to control MEFs, the cell spread area was smaller on 60 KPa compared to NT controls (Figure 4B). This was consistent with a previous study showing reduction in sensitivity of Sept-7 depleted cancer-associated fibroblasts to changes in ECM stiffness^32^.

**Figure 4.**
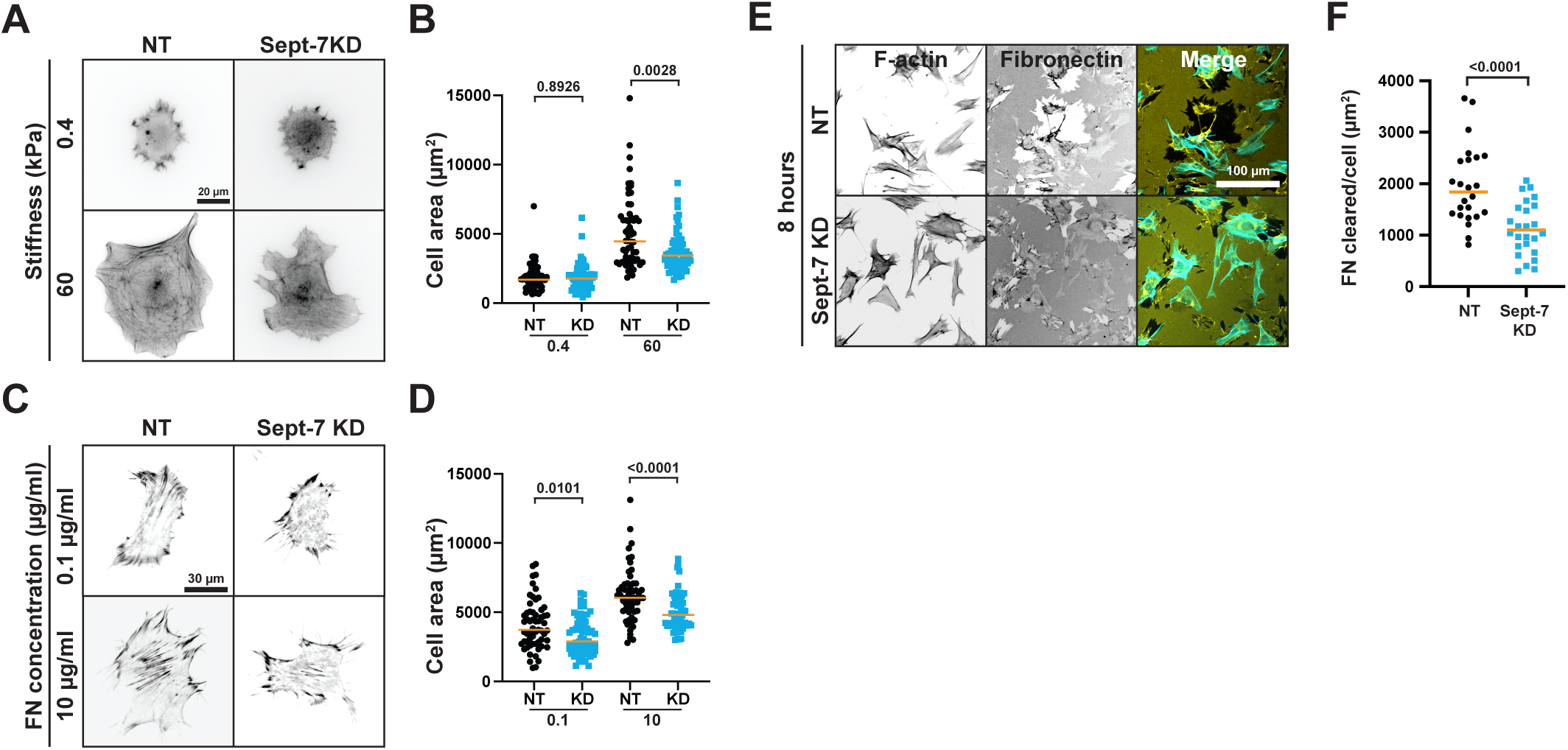
SEPT-7 enhances sensitivity of cells to changes in ECM cues and contributes to ECM remodelling. (A) Epifluorescence images showing NT and Sept-7 KD MEFs stained with F-actin and plated on soft (0.4KPa), and stiff (60KPa) polyacrylamide gels. (B) Quantification of cell area for NT and Sept-7 KD cells on soft vs stiff gels, n = 59 – 61 cells, with Kruskal-Wallis and Dunns multiple comparisons test. (C) TIRFM images of NT and Sept-7 KD of MEFs plated on low (0.1 µg/ml) and high (10 µg/ml) FN concentrations and co-stained with F-actin. (D) Quantification of cell area for NT and Sept-7 KD MEFs on low vs high FN concentrations, n = 59 – 61 cells, with Kruskal-Wallis and Dunns multiple comparisons test. (E) Epifluorescence images showing NT and Sept-7 KD MEFs plated for 8 hr on 10 µg/ml FN and co-stained with F-actin (cyan), and FN (yellow). (F) Quantification of the average area of FN clearance per cell in NT and Sept-7 KD cells, n = 24 – 25 images, with Mann-Whitney test.

Since our results show that formation of higher-order Sept-7 bundles and its regulation of perinuclear FAs is dependent on ECM binding, we next tested if similar to sensitivity to ECM rigidity, Sept-7 was critical in allowing cells to respond to changes in ECM density or haptosensing^33,34^. We plated NT control and Sept-7 KD MEFs on dishes either coated with 0.1µg/ml FN which reduces Sept-7 bundle formation at perinuclear FAs and elongation in peripheral FAs or on 10 µg/ml FN and stained the cells for paxillin and F-actin 4 hours after plating (Figure 4C). Imaging for cell-spread area showed that both NT control and Sept-7 KD MEFs failed to spread properly on 0.1µg/ml FN while increasing the FN density to 10µg/ml FN led to increased spreading and formation of stress fibers across the cell in the NT control and increased spreading with a reduction in perinuclear FAs and organised stress fibers in the Sept-7 KD cells (Figure 4C). Quantification of cell area confirmed that Sept-7 KD MEFs were significantly smaller than NT MEFs on both low and high FN densities (Figure 4D). Thus, like ECM stiffness sensing, Sept-7 increases sensitivity of cellular responses to changes in ECM density.

Lastly, based on the role of Sept-7 in formation of perinuclear FAs (Figure 3) and the known perinuclear localization of fibrillar FAs^35,36^, we asked if Sept-7 plays a role in ECM remodelling through regulation of perinuclear FAs. To test this, we first plated NT control and Sept-7 KD MEFs on 10 µg/ml FN and then fixed and immunostained the cells for FN and F-actin 8 hours after plating (to allow for ECM clearing in 2D). We then used epifluorescence microscopy to quantify the area of cleared FN at 8 hrs (Figure 4E). We observed areas of FN clearance in both conditions but the quantification of the average area of FN cleared per cell after 8 hours revealed a significant drop in ECM clearance in Sept-7 KD cells (Figure 4F). To test if this loss of FN remodelling was due to impeded cell migration or the ability of cells to remodel bound ECM, we tracked the migration of NT and Sept-7 KD cells labelled with SiR-DNA over 12 hrs. Quantification of cell migration speed revealed a slight reduction in migration speed in Sept-7 KD cells compared to NT control cells and no change in persistence or forward progress due to loss of Sept-7 (Figure S3E). However, the differences in cell migration speed were insufficient to account for differences in cleared FN between control and Sept-7 KD cells. This suggests that Sept-7 promotes the ability of fibroblasts to remodel the ECM by promoting remodelling and cell migration.

Taken together, these data shows that ECM and FA dependent localization of Sept-7 is critical in enhancing sensitivity of fibroblasts to sense changes in ECM cues and promotes the cells’ ability to respond and remodel the ECM.

## Discussion

Our results here show that Sept-7 is an FA protein that gets assembled into different higher order structures in FAs in an ECM dependent manner in fibroblasts. In addition to differences in their architecture, these structures also differentially localize to FA subpopulations with bundles of Sept-7 localizing to perinuclear FAs while elongated FA-like Sept-7 structures localize to peripheral FAs. Besides localizing to FA subpopulations, our results show an important role for Sept-7 in regulating these subpopulations. Most significantly in perinuclear FAs, Sept-7 not only increases maturation rate but also contributes to the stabilization and growth of these FAs. Functionally, we find that downregulation of Sept-7 expression results in fibroblasts losing their sensitivity to changes in ECM cues including stiffness and density as well as their ability to physically remodel the ECM. Collectively, these results show that Sept-7 is an important protein of the FA adhesome that via assembly of higher order structures determines the sensitivity of fibroblasts to sense changes to their ECM environment and regulate their functional response to changes in ECM cues.

Previous studies on the role of Sept-7 in regulating FAs have attributed this role of Sept-7 to its ability to interact and regulate F-actin outside and in the vicinity of FAs^37,38^. A recent study showed that septins can also target non-centrosomal microtubules to FA sites to drive FA disassembly^39^. However, as mentioned earlier, proteomic studies find several septin isoforms as part of integrin-based adhesion complexes or the adhesome^13^, suggesting a more direct role for septins in FAs. Here, our data using adhesion isolation and TIRF imaging confirms that Sept-7 is in FAs. While the specific mechanisms of FA recruitment are not known, a recent study using pull-down and mass spectrometry identified talin as one possible binding partner^21^. Talin is a large multi-domain FA protein that links the cytoplasmic tails of integrins to F-actin. Under mechanical forces, talin opens up to reveal a large number of binding sites for other FA proteins^40^. Investigating whether Sept-7 is one such FA protein will be subject of further studies. Other potential binding partners for septins include vinculin, LM07 and ZNF185 which were identified in a separate proteomics study investigating the Sept-9 interactome in human fibroblasts ^41^. Interestingly, since vinculin binding to talin is a tuneable mechanism required for ECM sensing, whether septins directly modulates vinculin-talin binding and thus enhances ECM sensitivity as we find here, should also be investigated further.

Our results here also lead to new questions about assembly of higher order septin structures. Septins form filaments by annealing hetero-oligomers which further form higher-order structures such as bundles and rings either through end-to-end binding or through lateral stacking^42^. Interestingly, Sept-7 is a component that is present in both fundamental units of septin oligomers, hexamers, and octamers and thus are an integral part of all higher order septin structures in a cell^43,44^. Our data on the ventral cell surface shows Sept-7 in 2 different structural forms, elongated FA-like in the peripheral FAs and longer bundles in the perinuclear region associated with perinuclear FAs. We show here that these structures while being dependent on the ECM have different reliance and sensitivity to integrin activation and ECM binding. The different form and mechanism of formation thus suggests distinct but ECM-dependent mechanisms of regulation of septin architecture on the ventral cell surface. Previous studies have shown that perinuclear septin bundles interact with perinuclear ventral actin stress fibers via Cdc42EP3 that stabilizes F-actin^19,45^. However, the fact that septin bundles are spatially confined to the perinuclear ventral surface and excluded from peripheral regions suggests additionally players in this mechanism. In addition, our results show that Sept-7 bundles specifically target ventral actin stress fibers and seems to have no effect on the actin cap on top of the nucleus even though Sept-7 can localize there. Along with the role of ECM binding in this process, this suggests that the alternate mechanisms are specific to perinuclear FAs that regulate the building of Sept-7 bundles. Even lesser is known about the regulation of FA-like peripheral Sept-7 structures. Their dependency on complete integrin activation and associated contractility suggests that these structures are directly or indirectly dependent on binding to FA proteins that undergo conformational changes at high forces or are associated with highly mature FAs. Here again, binding to talin or vinculin can provide a potential mechanism since if there are multiple binding sites for Sept-7 on talin or Sept-7 associates with multiple vinculins bound to talin, this can result in formation of FA-like structures when talin is elongated.

This study shows that loss of Sept-7 and the resultant loss in perinuclear FA function results in diminished ability of fibroblasts to sense and respond to biophysical changes in the ECM. A number of different mechanisms have been suggested by which cells tune their sensitivity and specificity to changes in ECM cues^46^. Our data here along with previous studies show that part of this mechanism relies on FA heterogeneity^47–49^. While FAs often look the same under the microscope when imaged with several canonical FA proteins, it is becoming clear that FAs within the cell are different from each other. However, very little is known about the specific roles of FA subpopulations in addition to their compositional and organizational differences. Our data shows that Sept-7 specifically plays a critical role in regulating perinuclear FAs and this correlates with the loss of sensitivity to ECM cues. Since the Sept-7 structures that coincides with this subpopulation are the perinuclear bundles, this suggests an overall organizational difference between perinuclear and peripheral FAs. It is also most likely that perinuclear FAs in our system are not one population but a few different populations of FAs with more specific functions that are currently unknown. Due to the effect of these perinuclear FAs on ECM remodelling, it is tempting to speculate that these perinuclear FAs are fibrillar adhesions that originate at the medial margins of classic FAs, changing protein composition and moving to the centre of the cell^50^. Paxillin however is thought to excluded from fibrillar adhesions, which instead contain tensin-1 that links F-actin to the cytoplasmic tail of integrins^36,51^. This shows that while the perinuclear FAs in our study are not fibrillar adhesions, there are some functional overlaps that needs to be investigated further.

In conclusion our work proposes spatially and mechanistically distinct roles for Sept-7 in the dynamics and function of perinuclear and peripheral FA subpopulations. Furthermore, we propose that Sept-7 is a critical FA component that promotes ECM mechanosensing and regulates the ability of fibroblasts to remodel the ECM.

## Methods

### Cell culture

Mouse embryonic fibroblasts were grown in Dulbecco’s modified eagle medium (DMEM + glutaMAX, Gibco, 61965026), 10% foetal bovine serum heat activated (Gibco, 10270106) and Penicillin/Streptomycin (Gibco, 15140122). Cells were kept at 37^0^C and 5% CO_2_.

### Magnetic bead-based FA isolation assay

Beads were functionalized with FN following previously described protocols^24,26^. Cells were cultured to 80 % confluency, washed with PBS, and incubated for 30 min in serum free media (SFM). FN coated beads were added to the cells for 30 min in SFM. Cells were lysed with NP-40 lysis buffer and lysates and bead fractions were collected and treated for 10 min with Benzonase (Sigma, E1014-25KU) to break up DNA/RNA. Bead fractions were then separated from lysates using a DynaMag-2 magnetic separator (Invitrogen, 12321D) and washed 3 times in NP-40 ready for Western blotting.

### Western blotting

Samples were resuspended in PBS and denatured with 4 x Laemmli sample buffer (Bio-Rad, 1610747) and boiled at 95^0^C for 10 min. Samples were treated with SDS-PAGE on a 4-12% Tris-Glycine gel (Invitrogen, XP041025BOX) and transferred to PVDF membrane (Biorad, 10026934) using Pierce transfer buffer (Invitrogen, PB7100). Membranes were blocked for 1 h in 3 % BSA (Sigma-Aldrich, A7906-100G) in TBS-T at room temperature. Primary Ab coupling was performed overnight at 4^0^C in 3 % BSA in TBS-T. Primary Abs used were: anti-mouse: Paxillin 1/3000 (BD Bioscience, 610052), Vinculin 1/800 (Sigma-Aldrich, V4505), focal adhesion kinase (FAK) 1/800 (Millipore, 06-534), anti-rabbit: Phospho-FAK 1/800 (Thermo Fisher, 44624G), phospho-paxillin 1/800 (Thermo scientific, 44-722G), Sept-7 1/800 (Thermo scientific, PA5-54755), GAPDH 1/5000 (Sigma-Aldrich, PLA0125). Membranes were washed in TBS-T 3 x 10 min before incubating for 1 h at room temperature (RT) with secondary Abs in 3 % BSA in TBS-T followed by 3 x 10 min wash in TBS-T and 1 x 5 min TBS. Secondary Abs used were: Starbright™ Blue 520 Goat Anti-Rabbit IgG (Biorad, 12005869), Goat anti-mouse IgG StarBright™ Blue 700 (Biorad, 12004158). All imaging was performed using a Bio-Rad ChemiDoc MP system.

### Immunostaining sample prep

Glass bottom dishes were coated with 10 µg/ml FN overnight in the fridge. Cells were plated for 4 hr at 37^°^C in DMEM before fixing for staining and imaging. For ECM ligand concentration experiments glass bottom dishes were coated with 0.1 or 10 µg/ml FN for 1 hr at 37^°^C, and then blocked in the fridge overnight in 2% BSA in TBS-T. For ECM stiffness experiments, polyacrylamide gels on 18 mm coverslips with a Young’s modulus of 0.4 or 60 Kpa were prepared and functionalized following a previously described protocol^52^ and coated with 10 µg/ml FN.

### Integrin activation assay

Glass bottom dishes were coated with 0.1 or 10 µg/ml FN in PBS, or PLL, for 1 hr at 37^0^C. Dishes were then rinsed with PBS and left overnight in 2% BSA in TBS-T and then rinsed with PBS. Cells were preincubated with 1 mM MnCl before plating on FN coated dishes for 4 hr at 37^°^C in DMEM and then fixed for staining and imaging

### Transfection

To analyze FA and Sept-7 dynamics, cells were transfected with 5 µg or 2,5 µg (single or co-transfection respectively) paxillin-mCherry, and/or Sept-7-YFP using Lipofectamine™ 3000 Transfection Reagent (ThermoFisher, L3000001). Cells were used 48 hr after transfection.

For Sept-7 knockdown experiments cells were transfected using with 20pM non-targeting (Dharmacon, D-001810-10-05), or Sept-7 targeting (Dharmacon, L-042160-01-005) siRNA. Cells were used 48 hr after transfection. For co-transfection with siRNA and plasmid, 5 µg paxillin-mCherry was transfected 24hr after siRNA transfection. Cells were used for experiments 24hr later.

### Immunostaining

Cells were fixed in 4 % paraformaldehyde (Thermo scientific, 28906) in cytoskeleton buffer (CB) for 20 min at 37^0^C. Cells were permeabilised in 0.5 % Triton-x (Alfa Aesar, A16046) in CB for 5 min, washed with 0.1 M Glycine (Sigma, 50046-250G) in CB for 10 min and then washed 3 times with TBS, all at RT, and then blocked with 2% BSA in TBS-T for 1 hr. Incubation of Primary Abs: Paxillin 1/400 (BD Bioscience, 610052), Sept7 1/400 (Thermo scientific, PA5-54755), FN 1/400 (Sigma-Aldrich, F3648-100UL) in 2% BSA in TBS-T was performed overnight at 4^0^C. Subsequent washes and incubations were in TBS-T. Cells were washed 3 x 5 min, incubated with secondary Ab: goat anti-mouse IgG 647 nm 1/400 (invitrogen, A1101), goat anti-rabbit IgG 568 nm 1/400 (Invitrogen, A11010) and phalodin 488 1/400 (Invitrogen, A12380) for 1 h in the dark at RT and then washed 3 x 5 min. Gels were mounted on slides with mounting media (Thermo scientific, P36980).

### Total internal reflection microscopy (TIRFM)

Images were acquired using total internal reflection microscope on a Nikon Eclipse Ti microscope with TIRF APO 100x 1.49 N.A. objective. Laser lines used were 488, 561, and 647 nm and emission and excitation filters were: GFP (mirror: 498–537 nm and 565–624 nm; excitation: 450– 490 nm and 545–555 nm; emission: peak 525 nm, range 30 nm) and mCherry (mirror: 430–470, 501–539, and 567–627 nm; excitation: 395–415, 475–495, and 540–560 nm; emission: peak 605 nm, range 15 nm), or Continuous STORM (mirror: 420–481, 497–553, 575–628, and 667–792 nm; excitation: 387–417, 483–494, 557–570, and 636–661 nm; emission: 422–478, 502–549, 581–625, and 674–786 nm). Images were acquired using a Teledyne Photometrics 95B 22 mm camera. For live cell imaging cells were kept at 37^0^C and images were captured every 10 – 30 sec over a 20 min timeframe.

### Confocal microscopy

Images were aquired using a Nikon Confocal A1RHD microscope with 488-, 561-, and 640-nm laser lines for F-actin, Sept-7, and paxillin respectively using a 60x Apochromal oil objective (N.A: 1.42).

### Structured illumination microscopy (SIM)

Image acquisition was performed using a Nikon N-SIM microscope with an LU-NV laser, and a CFR SR HP apochromat TIRF 100x oil objective (N.A: 1.49), 488 and 568 laser lines were used for F-actin Sept-7 respectively. An ORCA-flash 4.0 sCMOS camera (Hamamatsu Photonics K.K) was used and the images were reconstructed using in-built Nikon SIM software on NIS elements AR (NIS-A 6D and N-SIM analysis).

### Epifluorescence imaging

Image acquisition was performed using a Nikon Eclipse Ti microscope with an APO 20x 0.75 N.A. objective. Excitation and emission light was passed through a FITC (Exc. 457-487nm, Em. 503-538nm) or Cy5 (Exc. 590-645nm, Em. 659-736 nm) Semrock filter cube. Images were acquired on a Nikon DS-Qi2 CMOS camera. For live cell imaging, an environmental chamber (Okolab) was used to keep samples in a humidified 37^°^C and 5 % CO_2_ atmosphere. Cells were imaged every 5 min for 12 hr.

### Image analysis

*(All performed in Fiji unless stated otherwise)*

#### Cell area

All cell area analysis was performed on 20x epifluorescence images using manually created pipeline written in Julia (version 1.6), where Otsu thresholding was used to attain cell size.

#### Colocalization plot

(Figure 1C) were acquired by creating line plots of the average intensity values along cross sections of perinuclear or peripheral FAs of background subtracted images. Data was normalized using 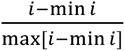 where i denotes intensity.

#### FA morphology

ROIs of perinuclear and peripheral FAs from background subtracted 100x TIRFM images were created. A median filter and Otsu thresholding was applied, and a mask created. FA sizes and numbers were then calculated using the built-in function to analyze particles. FA sizes were filtered to encompass a range of 0.20 µm^2^ < FA < 20 µm^2^.

#### FA dynamics

Kymographs were created of perinuclear FA sites from background subtracted 100x live cell movies, which were then used to measure minimum FA lifetimes and formation rates.

#### ECM remodeling

A manual threshold was used on fibronectin labeled epifluorescence 20x images to segment areas of the coverslips that were clear of fibronectin signal. A mask was created, and the segmented area calculated using the analyze particle’s function, which was then divided by the number of cells in the image to give an average area of FN cleared per cell.

#### Cell migration

Cell migration analysis was performed using the TrackMate plug-in^53^ using 20x epifluorescence timelapse images of cells labelled with the nuclear marker SiR-DNA (Tebubio, SC007). Cells were imaged over 12 hr and cells that were continually tracked for a minimum 8hrs were included in the analysis.

### Statistical analysis

All data was analysed using GraphPad Prism (version 10). Non-normally distributed data was analysed using a Mann-Whitney U test. Kruskal-Wallis test with Dunns post hoc was used for data with multiple comparisons. Dot plots or violin plots were used for data display, with orange horizontal lines showing medians.

## Acknowledgements

We thank Dr. Pontus Nordenfelt, Dr. Johan Malmström, Dr. Sebastian Wasserstrom and all the members of laboratory of cell and molecular mechanobiology (LCMM) for their discussion and support. The Sept-7 YFP plasmid was a kind gift from Dr. Helge Ewers at the Freie Universität Berlin. Johannes Kumra Ahnlide and Valeriia Grudtsyna are specially acknowledged for the help in developing code and maintaining image storage servers. Lund University Bioimaging Centre (LBIC) at Lund University is gratefully acknowledged for providing experimental resources. This research was funded by the Knut and Alice Wallenberg foundation (WS, VS), Wallenberg Centre for Molecular Medicine, Lund);Cancerfonden (VS, 19 0445 Pj and 22 2398 Pj Projekt grant) and The Royal Physiographic Society of Lund (WS, App: 43178).

## Supplementary Movie Legends

**Supplementary movie SM1**. TIRFM time-lapse imaging of MEFs transfected with Sept-7 YFP (grey) and mCherry Paxillin (magenta). Images taken every 10 seconds (elapsed time shown in min:s). Insets show perinuclear (second panel) and peripheral (right-most panel) regions of the cell. Montages in Figure 1D and 1F as well as line scans in 1E and 1G correspond to this movie.

**Supplementary movie SM2**. TIRFM time-lapse imaging of MEFs transfected with NT siRNA control for 48 hours and mCherry Paxillin (grey) prior to imaging. Images taken every 10 seconds (elapsed time shown in min:s). Inset shows perinuclear FA dynamics. Movie corresponds to Figure 3F (top panel). LUT inverted.

**Supplementary movie SM3**. TIRFM time-lapse imaging of MEFs transfected with Sept-7 targeting siRNA for 48 hours and mCherry Paxillin (grey) prior to imaging. Images taken every 10 seconds (elapsed time shown in min:s). Inset shows perinuclear FA dynamics. Movie corresponds to Figure 3F (bottom panel). LUT inverted.

---

**Figure S1.**
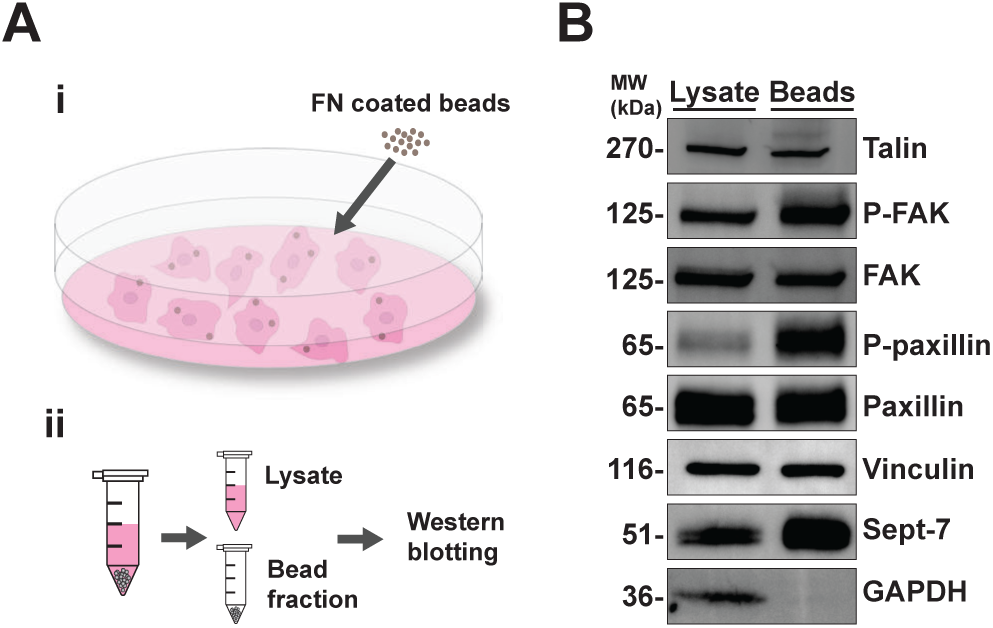
(A) Cartoon depicting magnetic bead-based FA isolation assay, (i) MEFs are cultured and incubated with 4.5 µm magnetic beads coated with FN. (ii) After cell lysis, the bead fraction is isolated using magnetic separation. Subsequently, Western blotting is employed to probe for focal adhesion (FA) associated proteins. Western Blotting images showing talin, focal adhesion kinase (FAK), paxillin and phosphorylated FAK, paxillin and phosphorylated paxillin, vinculin, Sept-7 and GAPDH. Note: Vinculin, Sept-7 and GAPDH were run on a separate gel due to close molecular weights with other proteins.

**Figure S2.**
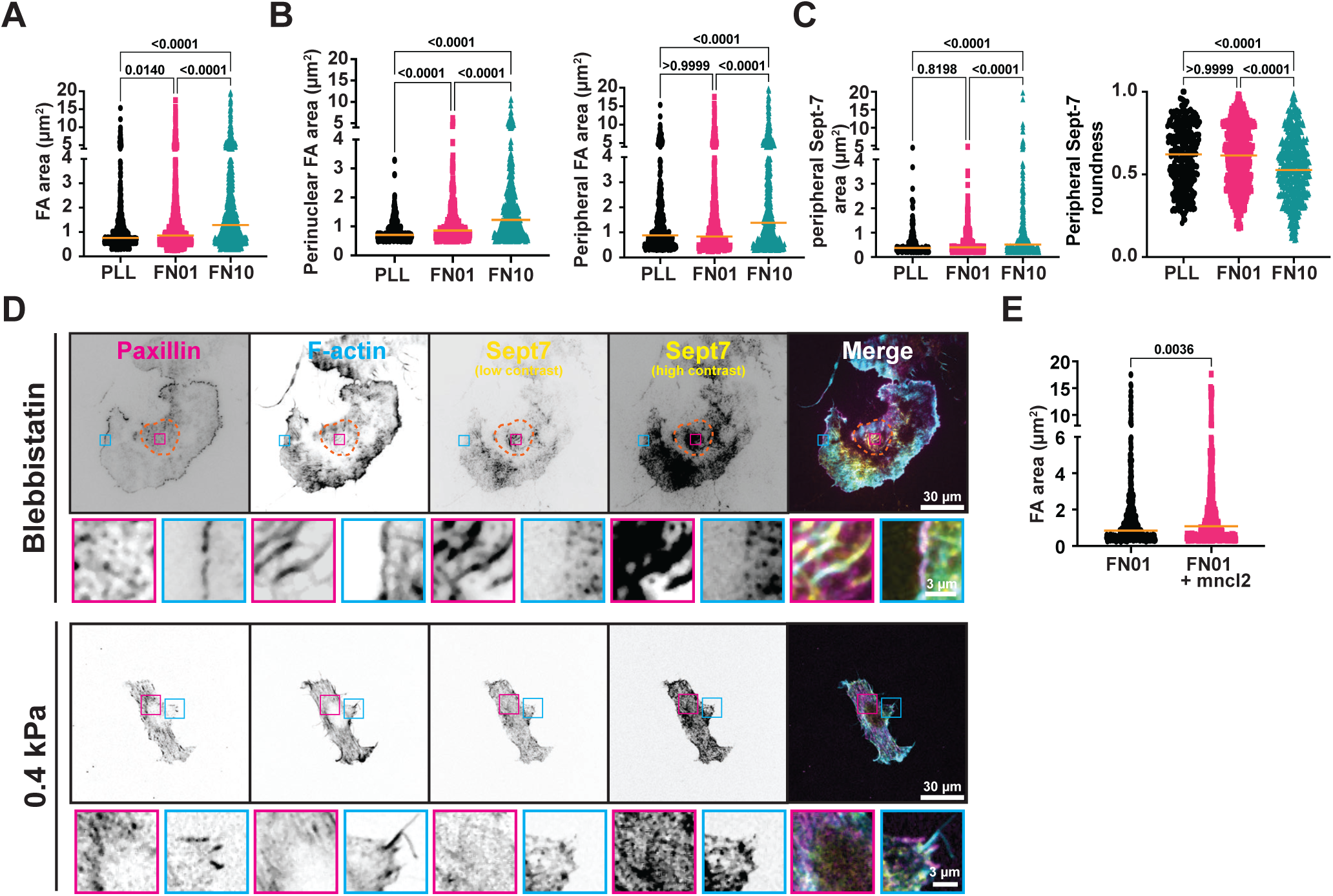
(A) Quantification of FA area of MEFs plated on PLL, 0.1 µg/ml FN, and 10 µg/ml FN, n = 860 – 1109 FAs. (B) Quantification of FA area of MEFs plated on PLL, 0.1 µg/ml FN, and 10 µg/ml FN, separated by perinuclear FAs (left plot, n = 396 – 546 FAs), and peripheral FAs (right plot, n = 434 – 560 FAs). (C) Quantification of peripheral Sept-7 puncta area (left plot), and roundness (right plot), for MEFs plated on PLL, 0.1 µg/ml FN, and 10 µg/ml FN, n = 152 – 399. (D) TIRFM images show paxillin (magenta), F-actin (cyan), Sept-7 (yellow, low, and high contrast), and merge of MEFs plated on 10 µg/ml FN and treated with blebbistatin (top panels), and 0.4 KPa gels coated with 10 µg/ml FN (lower panels), magenta and cyan insets highlight perinuclear and peripheral FAs respectively. (E) Quantification of peripheral FA area of MEFs plated on 0.1 µg/ml FN (data taken from S2B, right plot), and 0.1 µg/ml FN + 1mM MnCl2 n = 509 – 552 FAs. Kruskal-Wallis and Dunns comparisons test (A, B,C, and D), Mann-Whitney test (E).

**Figure S3.**
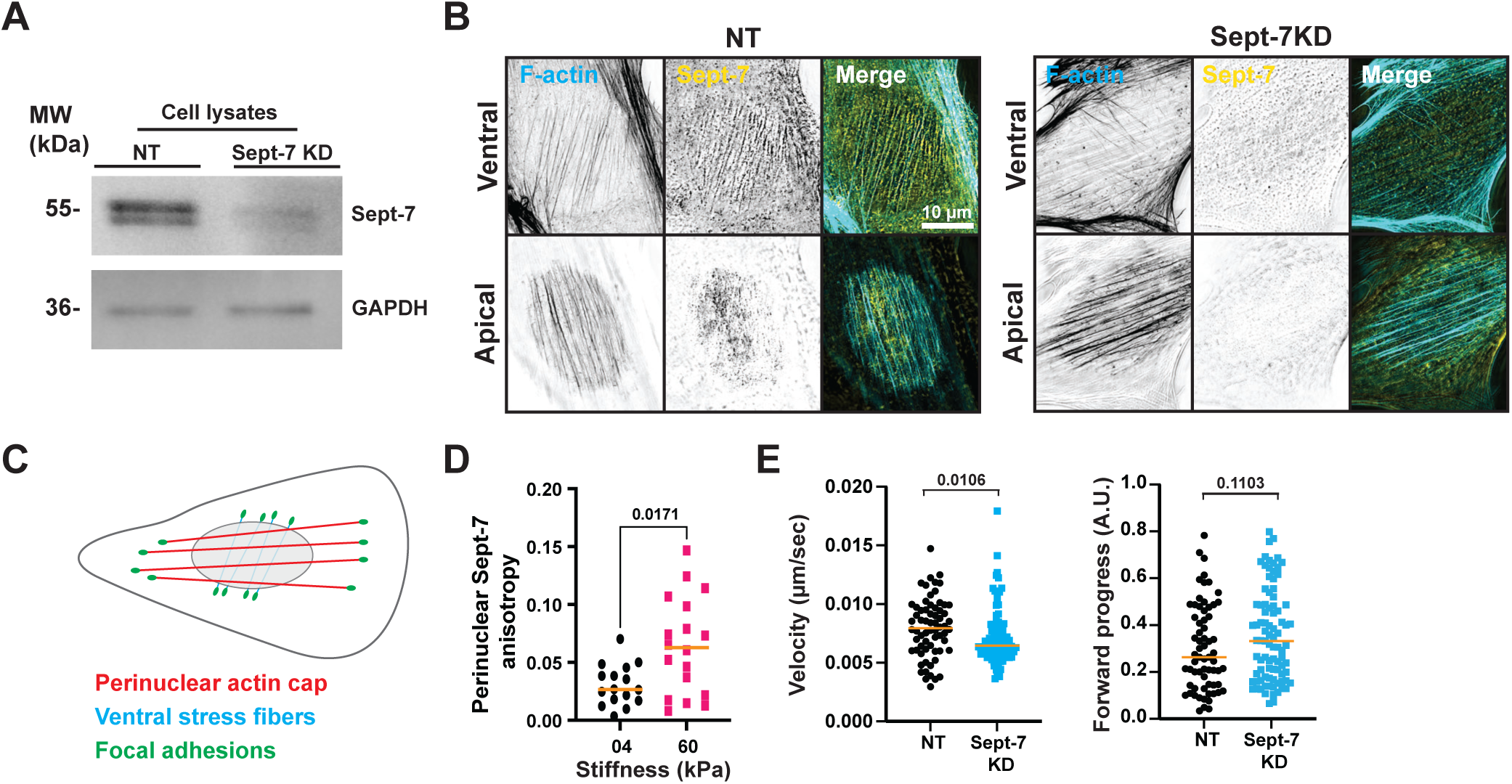
Representative Western Blotting images of cell lysates from MEFs showing Sept-7 and GAPDH after transfecting and incubating cells for 48 hours with NT or Sept-7 siRNA. (B) Representative 3D Structured Illumination Microscopy (SIM) images showing ventral SFs (ventral) and perinuclear actin cap (apical) of MEFs plated on 10 µg/ml FN treated with NT or Sept-7 siRNA and stained for F-actin (cyan), Sept-7 (yellow), and merge. (C) Cartoon depicting perinuclear actin cap apical to the nucleus, ventral SFs ventral to the nucleus, and FAs, of MEFs. (D) Quantification of perinuclear Sept-7 anisotropy of MEFs plated on soft vs stiff gels, n = 16 – 18 cells. (E) Quantification of cell velocity and forward progress of NT and Sept-7 KD cells imaged over 12 hr, n = 71 – 89 cells. Statistics taken using a Mann-Whitney test.

